# Investigating the Role of Management Decisions in Subspecies Hybridization Across the Wild Turkey’s Range

**DOI:** 10.1101/2024.11.07.622462

**Authors:** Amanda K. Beckman, Sarah A. Hamer, Gil G. Rosenthal, Leonard A. Brennan, Zach B. Hancock

**Affiliations:** Department of Biological Sciences Florida State University, Tallahassee, FL USA; Schubot Center for Avian Health Texas A&M University, College Station, TX USA; Ecology and Evolutionary Biology Interdisciplinary Program Texas A&M University, College Station, TX, USA; Veterinary Integrative Biosciences Department Texas A&M University, College Station, TX, USA; Department of Biology, Texas A&M University College Station, TX USA; Dipartimento di Biologia, Università degli Studi di Padova, Padova, Italy; Caesar Kleberg Wildlife Research Institute, Texas A&M University Kingsville, Kingsville, TX USA; Department of Ecology and Evolutionary Biology University of Michigan, Ann Arbor, MI USA

**Author notes:** **Correspondence:** Amanda K. Beckman, Florida State University Department of Biological Sciences 319 Stadium Dr. Tallahassee, FL 32304.

**Keywords:** Admixture, Conservation, Genomics, Introductions, *Meleagris gallopavo*, Translocations

## Abstract

The expanded geographic range and recovery to millions of wild turkeys across the country would not have been possible without management actions that included introducing and translocating individuals. However, the range-wide genetic impact of management decisions on one of North America’s greatest conservation success stories remains unknown despite the potential economic impact as hunters seek out easily identifiable subspecies for grand slams. In this study, we used DNA extracted from hunter-collected feathers from 29 states and Ontario to investigate genetic differences among turkeys in their historic and introduced ranges. Additionally, we compiled state-level management data to investigate how different management decisions are associated with the amount of admixture among subspecies. We found no difference in the amount of admixture in the turkey’s historic range compared to the introduced range. However, management decisions like as the number of subspecies introduced and the number of unique source states resulted in an increased level of admixture detected, but there was no relationship in admixture and the number of unique relocated counties. This first investigation into the hybridization among subspecies of wild turkey provides evidence that individual state’s management actions have influenced the genetic makeup of subspecies in that state.

## INTRODUCTION

Translocations and introductions were critical to the conservation success of wild turkeys (*Meleagris gallopavo)* in the last 100 years, but it is currently unknown how management has impacted the evolutionary trajectory of turkey subspecies within and outside their historical range. In their historic range, subspecies evolved and were separated geographically by known ecological barriers (Schorger 1966, Kennamer et al. 1992, Kuvlesky et al. 2020). The separation of eastern pine forests and the great plains prevented gene flow between eastern (*M.g. silvestris*) and Rio Grande subspecies (*M.g. intermedia*), while the preferred ponderosa pina habitat of Merriam’s (*M.g. merriami*) kept them separated from the Rio Grande’s western range. Gould’s (*M.g.* mexicana) prefer the more arid highlands of far southern New Mexico and Arizona and extending into Mexico which would prevent contact with Merriam’s on the northern edge of their range. These long-standing associations between turkey subspecies and specific geographic-based habitats have been confirmed with the calculation of divergence times between subspecies using modern and archaeological samples (Padilla-Jacobo et al. 2018).

The subspecies that experienced the most dramatic range expansion due to wild turkey conservation efforts was the Rio Grande subspecies (Eriksen et al. 2016, Chamberlain et al. 2022). Their historical range included Mexico, Texas, Oklahoma, Nebraska, Kansas, and South Dakota (Schorger, 1966), but due to management efforts they are now located in most states north and west of their historical range as well as Hawaii. The Merriam’s wild turkey also experienced human-influenced range expansion; their historical range was limited to Arizona, New Mexico, Utah, and Colorado (Schorger, 1966), but they have now been introduced north and west of their historic range (Chamberlain et al., 2022; Eriksen et al., 2016). In some western states, eastern wild turkeys, native to all the states west of the Rio Grande’s range, have been introduced as well. As a result, some states have two or more introduced subspecies without the wild turkey’s historic natural barriers to prevent hybridization. No studies have quantified the genetic impacts of wild turkey management on population structure across their entire range, but studies on a smaller scale have revealed that some populations have retained signatures of their source populations for decades (Rhodes et al. 1995, Mock et al. 2004, Latch and Rhodes 2005, Latch et al. 2006*b*, Seidel et al. 2013).

Population genomic tools provide a way to investigate the various evolutionary processes shaping population structure in wild turkeys, but historical management information can be used to provide additional context. Incorporation of genetics and management information have been previously used in population genetics studies of bobwhite quail (Williford et al. 2016) and white-tailed deer (DeYoung et al. 2003). Unlike the animal subjects of these previous studies, 90% of the domestic turkey (*M.g. domesticus*) genome has been assembled, and over 16,000 genes have been annotated (Dalloul et al. 2010); therefore, a wealth of genomic resources are available for turkeys that are not readily accessible for most wild species. From a conservation biology perspective, the rebound of wild turkey populations nationwide has been deemed one of the United States’ greatest wildlife management achievements. Quantifying how introduced populations have changed relative to their source populations and genetic differences between wild turkeys in their native versus introduced range has implications for improving management plans for other species.

In this study we investigated how levels of admixture varied between historic and introduced hybrid zones, and if levels of admixture were associated with states’ management decisions. Past studies quantifying hybridization among subspecies have focused exclusively on populations within the subspecies’ historic ranges (Latch et al. 2006*a*, *b*), this study is therefore first to assess subspecies hybridization across the wild turkey’s entire range. We predict that due to the lack of historic natural barriers, gene flow through secondary contact will be widespread throughout the western United States. Specifically, areas where multiple subspecies have been introduced will have widespread admixture detected. Additionally, we predict that as the number of subspecies introduced, unique introduction sources, and unique relocated sites each increase there will be increased admixture detected in that state.

## METHODS

### Sample collection

Samples were collected by recreational hunters, state and federal employees, and researchers from across North America. Volunteers were instructed to pluck approximately 10 breast feathers, making sure to keep intact the base of the feather that would be inside the skin of the bird (calamus). Volunteers provided their name, subspecies (henceforth called “hunter-identified subspecies”) and sex of turkey, date collected, state/county of collection, and source of collection (roadkill, hunter harvest, trapped, or other). Volunteers had the option of providing the following information: band number (if present), age class, city/ranch name, GPS coordinates, and habitat description of the collection site. Volunteers were instructed to store the samples in cool, dry, places in plastic bags or paper envelopes until they were shipped via standard shipping to College Station, TX. We used the geocode function in the R package “ggmap” to extract a GPS point in the county each sample was collected in to be used in further analyses (Kahle and Wickham 2013) and to determine whether the sampling location was within the wild turkey’s historic or introduced range (Schorger, 1966). Finally, we used the R package “fields” to calculate the great circle distance between all pairs of sampling coordinates (Douglas Nychka et al. 2021).

From the over 700 samples we received, we narrowed down the sample size to the 384 to match the probe design kit we purchased using a stratified random selection process for state, county, and submitter as follows: First, we divided the 384 by the number of states and Canadian provinces we collected samples from (30) to determine the value of 12-13 samples per state/province. Not all states/provinces had 12-13 samples, so recalculations were made to equally disperse the remaining slots in states with more than 12 samples. Within each of those states, we maximized geographic coverage by selecting samples from as many counties as possible. Finally, if more than once sample was selected from the same county we selected samples collected by different people.

Last, we wanted to collect more detailed information about turkey management across North America to further investigate how different management actions are associated with the levels of admixture detected. For all states where we collected samples, we contacted the appropriate wildlife official in each state asking for any turkey translocation/introduction records or information to compile into our own database.

### DNA extraction and sequencing

DNA was extracted from the calamus of the feathers; ∼1 cm from the base of the feather was cut into small pieces using sterile scissors, tweezers, and petri dishes. Approximately 4-6 feathers were used per individual to obtain the minimum target DNA concentration of 10 ng/μL suggested for effectively sequencing the probes. A basic salt DNA extraction was used; a lysis buffer and Proteinase K were added to 2 mL microcentrifuge tubes containing the cut-up pieces of calamus, and tubes were incubated at 55 C for 24-48 hours. Next, two ethanol washes were used to separate the DNA from other materials in the feather. Last, the tubes were left open on the bench overnight to allow any remaining ethanol to evaporate, and then the DNA pellet was dissolved in TE Buffer. After allowing the pellet at least 24 hours to precipitate into the buffer, sample concentrations were quantified using a Qubit 2.0 Fluorometer. Extracted DNA was stored at -20 C until all samples were submitted for sequencing. Extracted products were sent to the Texas A&M Institute for Genome Sciences and Society for library preparation and sequencing using the NovaSeq XP workflow.

Extracted DNA from plucked feathers are often subject to low to medium DNA degradation (Bush et al. 2011), but we used the Allegro targeted genotyping method which requires a low starting concentration of DNA and is suitable for small-medium sized fragments of DNA (NuGEN Technologies, San Carlos, CA, USA). Briefly, the DNA was fragmented using enzymes, adapters were ligated to the fragments, and then the fragments were extended and amplified. Samples were then pooled for library amplification and purification before being sequenced by the Nextseq 500 system, where each single nucleotide polymorphism (SNP) was sequenced to ∼100x.

SNPs were chosen from a previously published dataset containing 5 million SNPs found in turkeys (Aslam et al. 2012). 10,000 SNPs were chosen that were equally spaced throughout the genome, approximately every 100kb (kilobase). Choosing SNPs in this random way minimizes the effects of SNP ascertainment bias compared to using a previously developed SNP array (Geibel et al. 2021). Additionally, previous research showed that 10,000 SNPs will provide more than enough genetic information for a population or hybridization study (Turakulov and Easteal 2003, Morin et al. 2009). However, the genome version that was used previously to determine SNPs in turkeys (Turkey_2.01) was no longer the current genome version (Turkey_5.1;(Dalloul et al., 2010)). We converted the coordinates of the 10,000 selected SNPs from Turkey_2.01 to Turkey_5.1 using UCSC’s liftOver Linux executable (Kuhn et al. 2013).

### Bioinformatics

We worked with Interval Bio, a company specializing in bioinformatics and large-scale computation, to complete alignment, variant calling, and filtering. First, the Allegro probes were trimmed from the reads using Trimmomatic (Bolger et al. 2014). Then, reads were aligned to the reference genome using BWA (Li and Durbin 2009). FreeBayes was used to call genotypes at target positions and regions flanking the target that had a minimum of five reads and merged both targets and flanking SNPs into one VCF file (Garrison and Marth 2012). Finally, filtering to exclude indels/multi-allelic loci, variants on unplaced assemblies, loci with a depth less than 20, variants with minor allele frequency (MAF) less than 0.05, and loci missing data for more than 20% of samples was accomplished using BWA (Danecek et al. 2021). BWA was also used to remove samples that were missing more than 10% of remaining loci. Finally, pruning for SNPs in linkage disequilibrium (LD) was done using plink using the parameters “indep-pairwise 50 5 0.5” corresponding to calculating the pairwise genotypic correlation in 50 SNP windows, removing a pair of SNPs if the LD is greater than 0.5, and then moving the window 5 SNPs.

#### Analyses

We conducted the genetic differentiation and admixture analyses for this study using the R program “conStruct” (Bradburd et al. 2018). This program takes an allele frequency matrix, geographic distance matrix, and list of sample coordinates to categorize natural genotypic variation without any *a prioi* defined population structure. conStrct improved on previous algorithms for detecting population structure, like STRUCTURE, by incorporating isolation by distance using an explicit spatial component. conStruct uses a cluster-based approach to determine populations by minimizing disequilibrium for each cluster. ConStruct is also robust to situations where geographically separated populations interact, like post-range expansion (which would be similar to the situation of introduced populations retaining some signatures of their geographically distant source populations).

To determine the optimal number of clusters (K) for further conStruct analyses, we compared the layer contributions across different K values. The maximum number of clusters investigated was K=4, for the number of subspecies represented in the samples we collected. We used conStruct to match layers across runs and plotted the layer contributions to determine the point where layers were no longer explaining a meaningful amount of covariance. After the optimal K value was determined, we evaluated the diagnostic plots that conStruct outputs to evaluate model convergence. Last, we created the admixture graphs with samples oriented by longitude and latitude pie chart maps using conStruct.

From the admixture proportions output by conStruct, we calculated an admixture metric *m*_1_, given by equation 1, a heuristic measure representing an individual’s level of gene flow among subspecies:

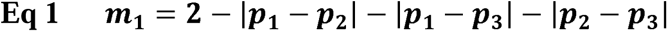

For each individual, a set of coefficients (*p*_1_, *p*_2_, *p*_3_) was calculated which represents the fractional classification into distinct groupings. By definition, *p*_1_ + *p*_2_ + *p*_3_ = 1. The values of m are between 0 and 2 (inclusive) where *m*=0 indicates no gene flow among subspecies (eg., *p*_1_ = 1, *p*_2_ = *p*_3_ = 0) and m= 2 represents an individual with equal ancestry across all three groupings (eg, *p*_1_ = *p*_2_ = *p*_3_).

We also calculated the admixture metric a second way that did not result in samples with admixture from three subspecies having a larger metric value than subspecies that were admixed between two subspecies. m_2_, calculated from equation 2 considers how much of the sample’s genome is admixed when the largest coefficient (*p*_mUX_) is removed:

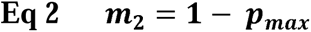

We evaluated how the historic/introduced state designation and hunter-identified subspecies predicted the admixture metric for each individual sample using a t-test and one-way analysis of variance (ANOVA) respectively. Next, we calculated the average and standard deviation of the admixture metric for all samples in each state. We used ggpmisc to fit and visualize linear models predicting statewide admixture from different management actions such as: number of unique subspecies translocated/introduced, number of unique source states, and number of unique relocated counties.

## RESULTS

We submitted 384 samples collected from across North America for sequencing (Figure III-1) and retained 367 individuals from 29 states and Ontario after filtering for missing data. After quality control filtering, the final number of shared filtered bi-allelic SNPs at target and flanking regions was 31,488, and after LD pruning the final number of shared SNPs across samples was 24,814.

We received replies from state officials and obtained management records from Ontario and 19 out of the 29 states we collected samples from. For states we did not hear back from, we were able to partially put together management information through information provided from other states and from previously published range maps (Chamberlain et al., 2022; Eriksen et al., 2016). We recorded the management documents provided by each state in our own dataset where each row in the dataset represented a unique year and source/relocated location. We did not include within-county movements. In total, we documented 3,800 unique turkey movement events.

By comparing models with different numbers of layers, we were able to determine the optimal K value for further analyses. The fourth layer only explains 1.9% of the observed variance, indicating K=3 is the optimal number of layers for our analyses. The clusters roughly correspond to Rio Grande, eastern, and Merriam’s/Gould’s. Next, we evaluated fit for the K=3 model by evaluating the trace plots of the Markov Chain Monte Carlo (MCMC) parameter estimates, which showed that the model had converged and did not need to be run for further iterations since the plots had plateaued. Additionally, the explained covariance for the observed geographic distances has good fit within the 95% confidence interval.

We plotted the admixture proportions as pie charts on a map to evaluate broad geographic trends of gene flow. When evaluating samples longitudinally, trends for all three clusters are observed (Figure 1). The admixture proportions of the westernmost samples in Hawaii and California closely resemble most samples collected in the middle of the United States. Also in the western US, there is a clear cluster of Merriam’s before the transition to samples being mostly comprised of Rio Grande genotypes. As expected, as samples move further eastward more samples with increased proportions of the eastern subspecies’ genotypes are observed.

**Figure 1.**
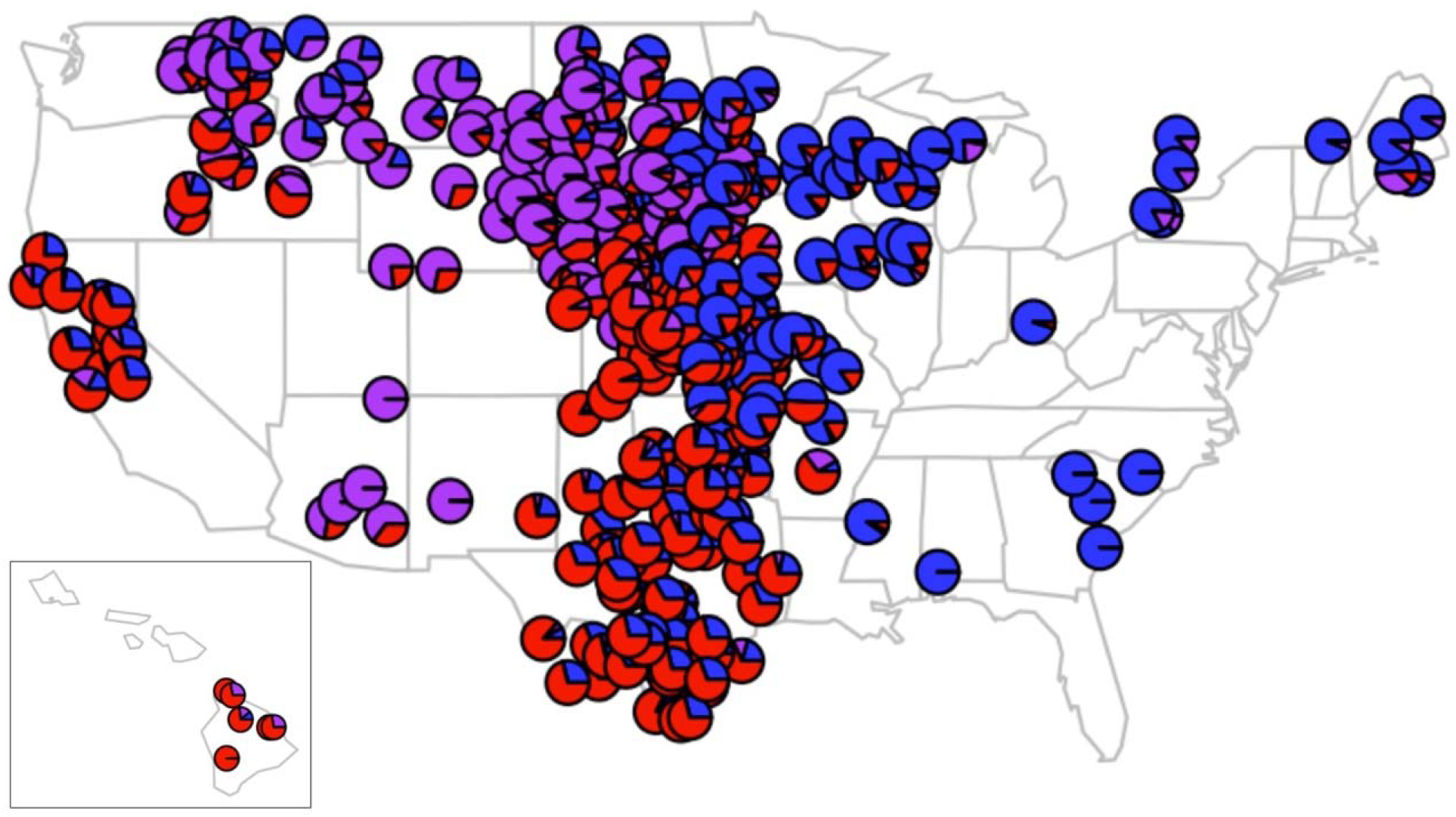
Pie charts showing admixture proportions of subspecies of wild turkeys across the US and Ontario. The sample coordinates used to produce this graph had random noise added to better visualize the pie charts. The colors correspond to eastern (blue), Rio Grande (red), and Merriam’s/Gould’s (purple) subspecies.

We summarized the individual admixture proportions calculated in conStruct into a single admixture metric that we measured two different ways. There was no relationship between the sample’s admixture metric and whether the sample was from the introduced or historic range of wild turkeys (*p_m1_ = 0.37, p_m2_ = 0.68*). However, several significant differences were observed in admixture metrics of different hunter-identified subspecies (Figure 2). Hunter-identified hybrids had a significantly higher admixture metric than all the other categories (*P < 0.001*), and the admixture metric for Rio Grande samples were significantly larger than eastern-identified samples (*P = 0.0169*) when the admixture metric was calculated using equation 1 (Figure 2A). Similar results were observed when the admixture metric was calculated equation 2 (*P = 0.0062*), with the addition of a significant relationship between the unknown category and the eastern subspecies (*P < 0.001*; Figure 2B).

**Figure 2.**
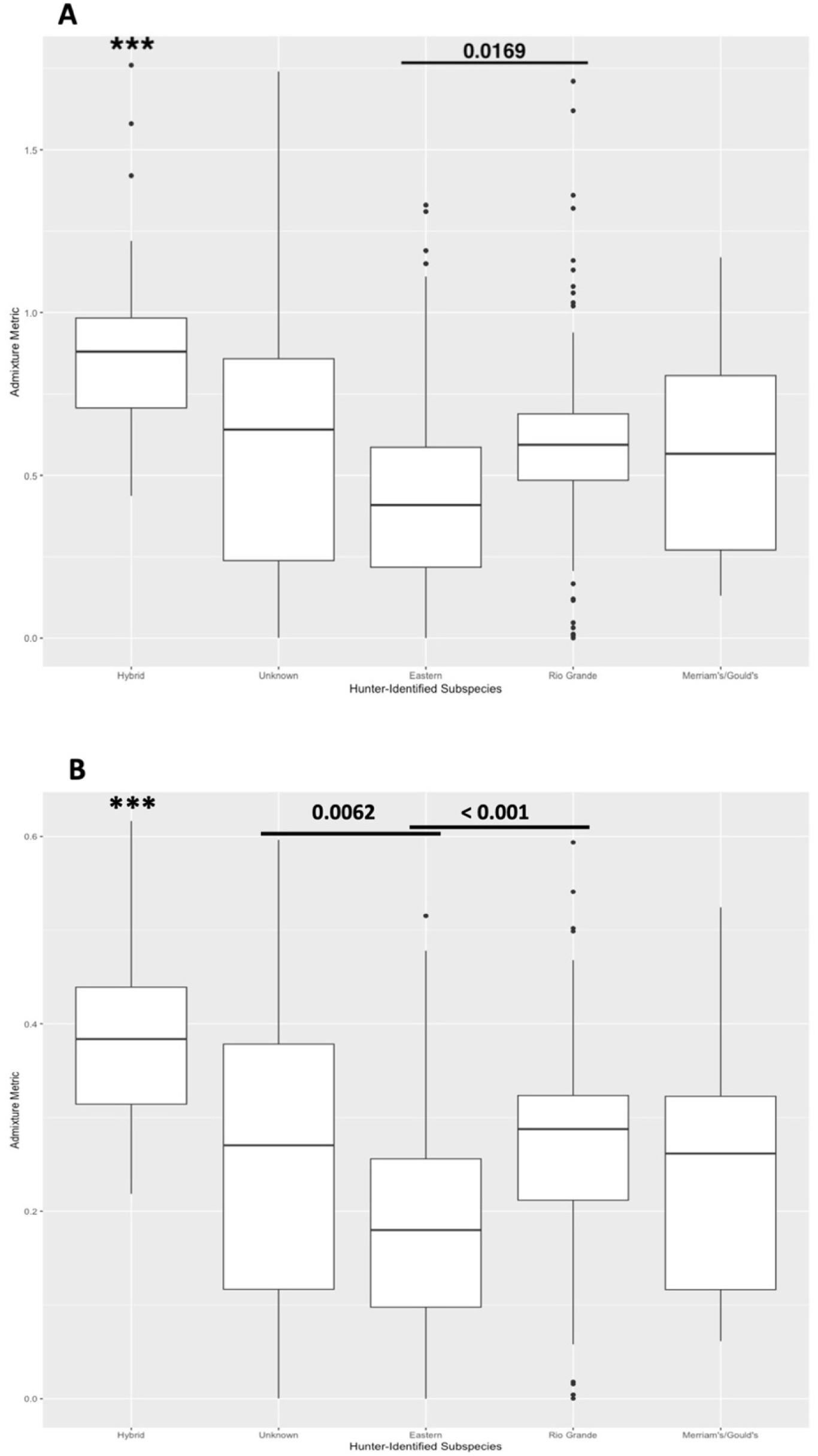
Individual admixture metrics compared across hunter-identified subspecies of wild turkeys. P-values above comparisons were calculated using a one-way ANOVA with a Tukey correction. Admixture metric was calculated in two different ways, m_1_ (A) and m_2_ (B). *** denotes a p-value < 0.001 against all other subspecies.

We took the average of all the samples in each state and evaluated whether the statewide admixture values were correlated with different turkey management actions. First, we determined whether the number of subspecies translocated and introduced influenced the average level of admixture in each state (Figure 3). The exact number of subspecies introduced in California is not firmly established (Gardner et al., 2004), so we made graphs where California was classified as having four subspecies introduced (Figure 3 A&C) and six subspecies introduced (Figure 3 B&D). Despite the number of subspecies introduced in California, there was a significant relationship between number of subspecies translocated/introduced and the average admixture metric for that state.

**Figure 3.**
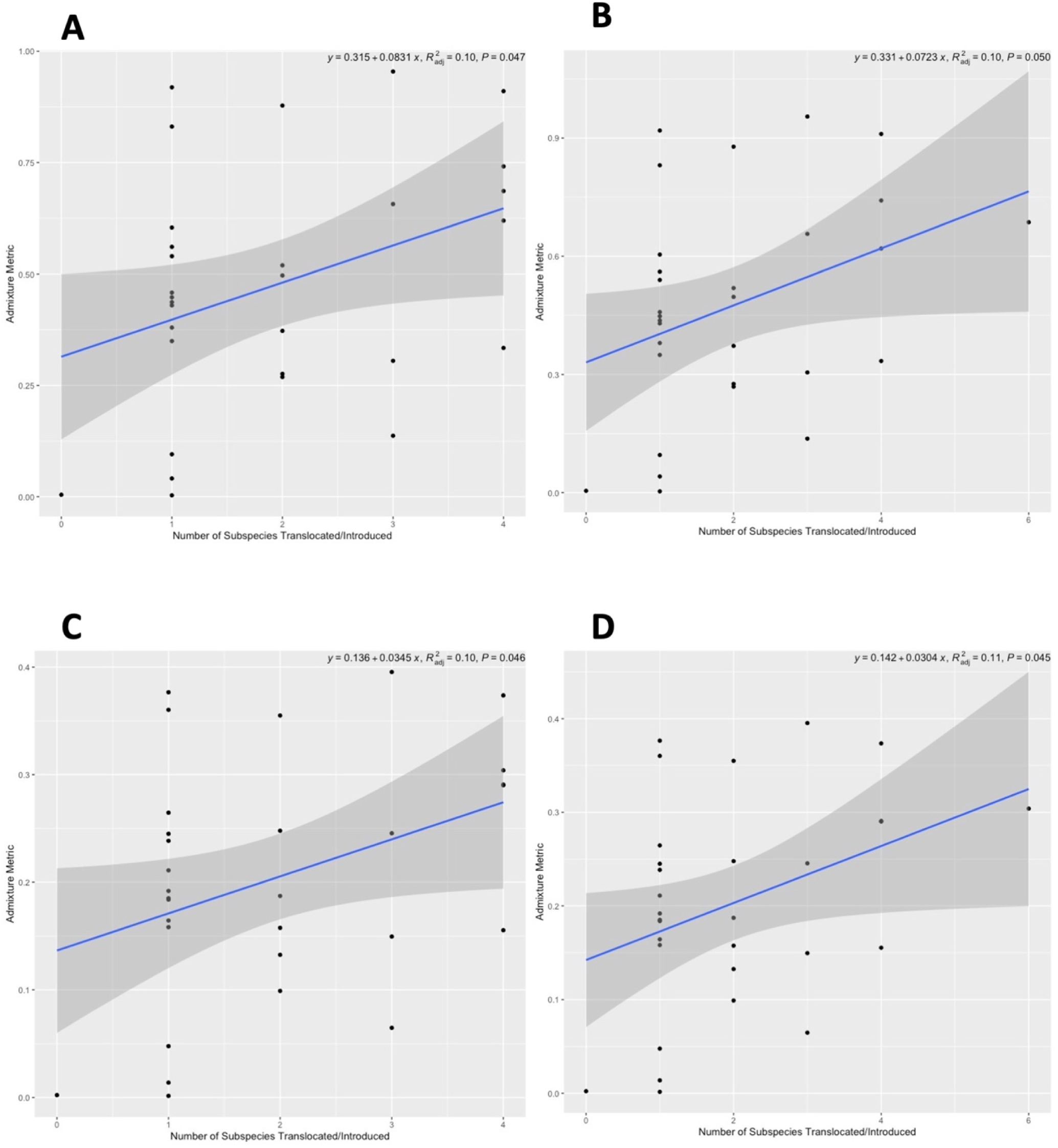
The average admixture metric for each state as predicted by the number of wild turkey subspecies that were translocated or introduced as part of management in that state. Unique introductions up for interpretation were done in California, so the total of number of introduced subspecies in California was plotted for four (A,C) and six (B,D) subspecies. The fitted linear model is added as a blue line on each graph. Model slope, fit, and p-value are provided at the top of the graphs. Admixture metric was calculated in two different ways, m_1_ (A,B) and m_2_ (C,D).

For the comparisons of average state admixture metrics and unique source states and relocated counties, we filtered the dataset to only include Ontario and the 19 states from which we directly received management information from so the results were not skewed by states that did not respond. As the number of states where turkeys were sourced from increased, the statewide average admixture metric significantly increased (Figure 4). However, the number of unique counties where turkeys were introduced to does not explain any variation in statewide average admixture metrics (Figure 5).

**Figure 4.**
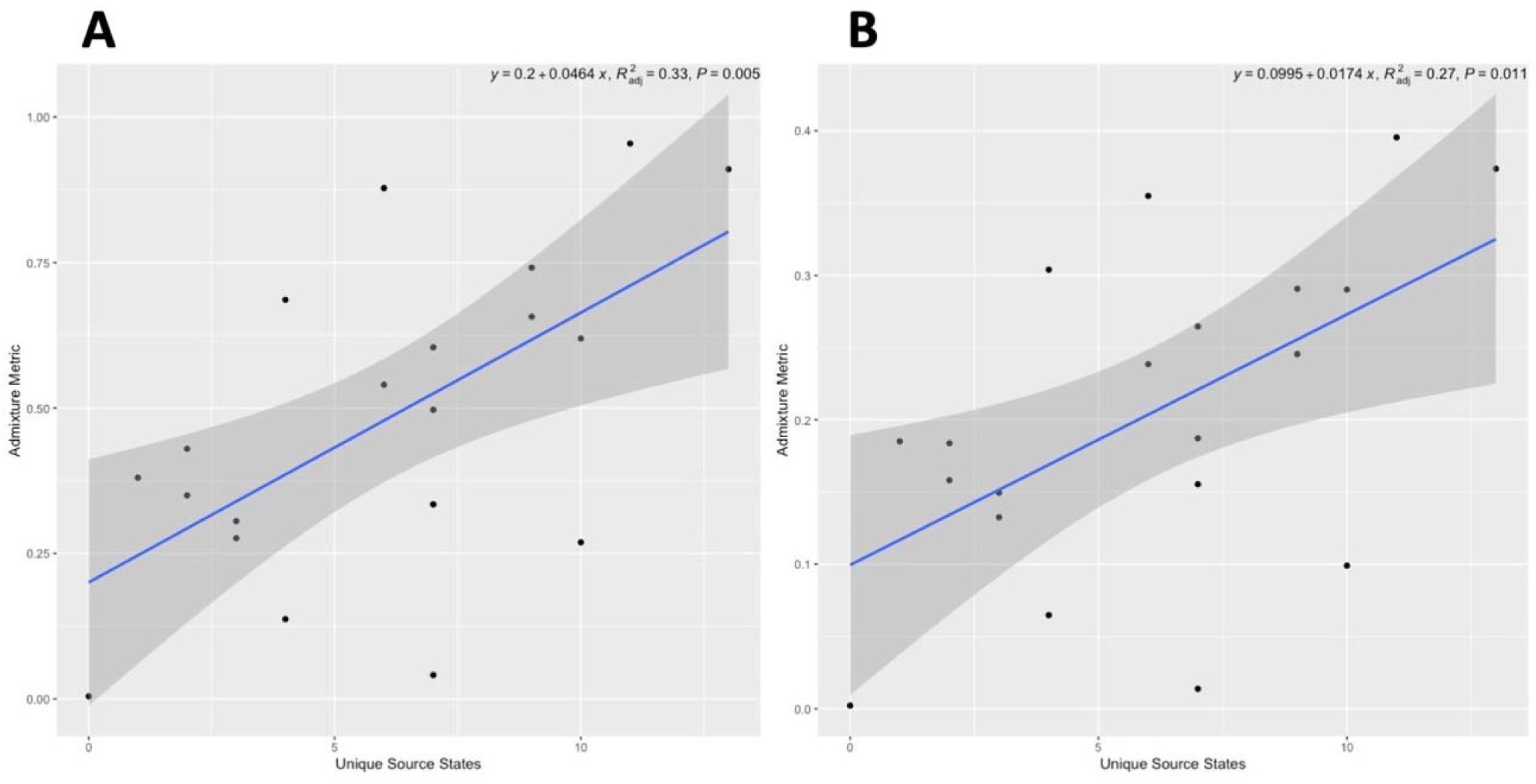
The average admixture metric for each state plotted by the number of unique source states that wild turkeys were obtained from for translocation/introductions. The fitted linear model is added as a blue line on each graph. Model slope, fit, and p-value are provided at the top of the graphs. Admixture metric was calculated in two different ways, m_1_ (A) and m_2_ (B).

**Figure 5.**
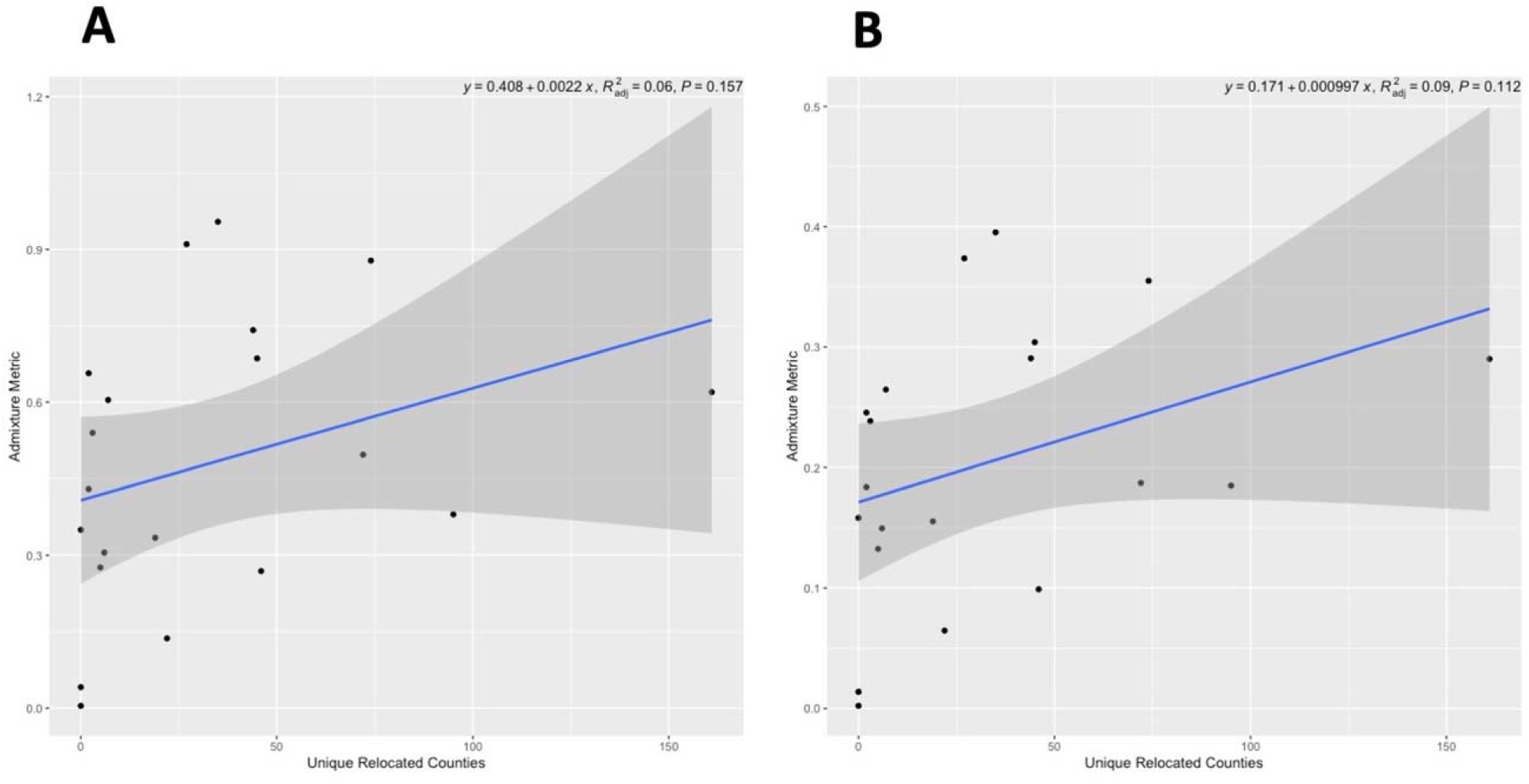
The average admixture metric for each state plotted by the number of unique counties where wild turkeys were relocated. The fitted linear model is added as a blue line on each graph. Model slope, fit, and p-value are provided at the top of the graphs. Admixture metric was calculated in two different ways, m_1_ (A) and m_2_ (B).

## DISCUSSION

We describe widespread admixture between subspecies of wild turkeys throughout North America, as one method toward quantifying gene flow among subspecies in historic hybrid zones and human-caused hybrid zones in the wild turkey’s introduced range. No difference in admixture among subspecies is observed in the wild turkey’s introduced and historic ranges. This is perhaps due to new barriers to hybridization the introduced turkeys are being separated by, or because the historic barriers to hybridization are eroding (deforestation of eastern pine forests) so admixture in the historic range is now indistinguishable from the introduced range where subspecies are readily breeding with one another.

Through the state-level translocation data that we compiled, we found that the number of subspecies introduced, and the unique source locations affected the level of admixture detected, but the number of unique source counties were not associated with the level of admixture detected. This indicates that inter-state decisions about the subspecies brought in and what states translocated/introduced individuals are from are associated with the level of admixture more than within-state decisions, which highlights the importance of forming wider geographic management networks instead of individual states making management decisions.

By comparing layer contributions across different values of K we were able to verify that K=3 was the optimal number of clusters for the dataset, validating that Merriam’s and Gould’s subspecies were clustered together in our analyses. Additionally, we looked at the results for the K=4 run and the samples in the extreme southwest US that were the hunter-identified Gould’s samples do not form the fourth cluster, which ended up being a minor genotype observed in the introduced northwestern US.

Geographic trends that have been well-described in the turkey literature for decades (Chamberlain et al., 2022; Kennamer et al., 1992; Schorger, 1966) are reflected in the admixture proportion graphs and map from our analyses. The historical west-east transition of Merriam’s and Gould’s in the mountainous regions of the southwest US, to Rio Grandes in the great plains, and then increasing proportions of eastern genotypes moving eastward from the historic hybrid zone with Rios. The westernmost samples in Hawaii and California closely resemble the admixture proportions of samples in the great plains which makes sense from a historical and climatic perspective since Hawaii and California both sourced mostly Rio Grandes for their introduction efforts and have warmer climates.

One limitation of the methods we used to calculate admixture is that the phylogenetic relationships of subspecies are not considered. For example, the eastern and Rio Grande subspecies share a more recent common ancestor relative to the other subspecies (Padilla-Jacobo et al. 2018). Without incorporating this into the model calculating admixture, some genotypes will erroneously be categorized as Rio Grande or eastern when they are really shared from their common ancestor. Future work could use models that incorporate phylogenetic methods (Hibbins and Hahn 2022) or using local ancestry inference to assign representative individuals from each cluster to compare the remaining samples to (Guan 2014).

Quantitatively summarizing the admixture proportions allowed to us further analyze how different designations or management actions affected the level of admixture detected. Hunter-identified hybrids had a significantly higher admixture metric than all the other categories, which indicates hunters can accurately identify hybrids phenotypically in the field. Additionally, since hybrids have a higher admixture metric than the unknown category, this provides evidence that hunters were accidentally not including subspecies information opposed to being unsure of what subspecies and not writing anything. Within-species feather color is known to be an incredibly variable trait, and when hybridization is present color can become even more variable (Ng and Li 2018). However, in this dataset most hunters correctly identified subspecies and hybrids, even in regions with multiple subspecies.

## CONSERVATION IMPLICATIONS

Without a doubt, wild turkey conservation efforts over the last hundred years have effectively saved the species. However, if the goal of turkey management is to preserve distinct subspecies of wild turkeys, then future management efforts may benefit from consideration of genetic data into management actions. Without alternative intervention, it is possible that recognizable subspecies could eventually deteriorate as admixture increases. Additionally, hybrid turkeys are already distinctly recognizable by turkey hunters, indicating that hybrids could eventually diverge into their own hybrid-origin subspecies. Aside from the biological implications of wanting to conserve subspecies, there potentially are very real economic implications if states only have recognizably hybrid subspecies and hunters continue to seek out opportunities to hunt subspecies they can clearly distinguish. Finally, further range-wide analyses and publication of open-access current and historic management wild turkey data should be prioritized to better inform management plans for currently imperiled species.

## ACKNOWLEDGEMENTS

The authors did not have any conflicts of interest relevant to this study. First, we would like to thank the hunters, state and federal employees, and researchers that donated their time and feathers, because without them this project would not have been possible. We also want to thank the state wildlife officials and NWTF employees that assisted us getting in contact with hunters, and those that provided state management information.

## ETHICS STATEMENT

Our study followed relevant regulations and guidelines and did not involve any handling of live animals. Authorization to salvage and collect feathers was provided through Texas Parks and Wildlife Department Scientific Permit No. SPR-0219-032.

